# Invalidity of light sensor data in field studies and a proposal of an algorithmic approach for detection and filtering of non-wear time

**DOI:** 10.1101/2021.08.11.455859

**Authors:** Larissa C. Hunt, Josef Fritz, Michael Herf, Lorna Herf, Celine Vetter

**Author notes:** Corresponding author: Dr. Céline Vetter; Department of Integrative Physiology, University of Colorado, Boulder. L.C.H. and J.F. contributed equally to this work. JF is now at the Department of Medical Statistics, Informatics and Health Economics, Medical University of Innsbruck, Innsbruck, Austria.

## Abstract

Wearable light sensors are increasingly used in intervention and population-based studies investigating the consequences of environmental light exposure on human physiology. An important step in such analyses is the reliable detection of non-wear time. We observed in light data that days with less wear-time also have lower variability in the light signal, and we sought to test if the standard deviation of the change between subsequent samples can detect this condition. In this study, we propose and validate an easy-to-implement algorithm designed to discriminate between days with a non-wear time >4h (“invalid days”) vs. ≤4h (“valid days”) and investigate to which extent values of commonly used physiologically meaningful light variables differ between invalid days, valid days, and algorithm-selected non-wear days. We used 83 days of light data from a field study with high participant compliance, complemented by 47 days of light data where free-living individuals were instructed not to wear the sensor for varying amounts of time. Light data were recorded every two minutes using the pendant-worn *f.luxomete*r light sensor; validity was derived from daily logs where participants recorded all non-wear time. The algorithm-derived score discriminated well between valid and invalid days (area under the curve (AUC): 0.77, 95% CI: 0.67-0.87). The best cut-off value (*i.e.*, highest Youden index) correctly recognized valid days with a probability of 87% (“sensitivity”), and invalid days with a probability of 63% (“specificity”). Values of various light variables derived from algorithm-selected days only (median: 264.3 (Q1: 153.6, Q3: 420.0) for 24h light intensity (in lux); 496.0 (404.0, 582.0) for time spent above 50-lux) gave values close to those derived from self-reported valid days only. However, these values did not significantly differ when derived across all days compared to self-reported valid days. Our results suggest that our proposed algorithm discriminates well between valid and invalid days. However, in high compliance cohorts, distortions in aggregated light measures of individual-level environmental light recordings across days appear to be small, making the application of our algorithm optional, but not necessary.

## INTRODUCTION

Light is the primary zeitgeber for the human circadian system (Duffy and Wright, 2005; Roenneberg *et al.*, 2013). Objective measurement of light using wearable sensors from various providers is increasingly used to study the consequences of light exposure patterns on human physiology, behavior, and health in the field. For example, Wams et al. demonstrated that the quality and architecture of sleep is associated with preceding light exposure after equipping 20 healthy participants under ambulatory field conditions for six days with the wrist-worn MotionWatch 8™ (MW8™, CamNTech Ltd., UK) light intensity monitoring device (Wams *et al.*, 2017). Obayashi et al. observed that brighter evening and nighttime light intensities were associated with longer sleep onset latency using the Actiwatch 2 device (Respironics Inc., Murrysville, PA) in 192 Japanese elderly individuals (Obayashi *et al.*, 2014). Additionally, in field studies about night and shift work, a partly light-mediated effect of night work on melatonin production was demonstrated (Papantoniou *et al.*, 2014; Daugaard *et al.*, 2017; Razavi *et al.*, 2019) using the HOBOware light intensity data logger (Onset Computer Corporation) in the first, the Philips Respironics Actiwatch Spectrum in the second, and the Daysimeter - a small, self-contained, battery-operated headset - in the third study. It was also shown that timing and intensity of light correlate with body mass index using 7 days of light data of the wrist-worn AW-L Actiwatch (Mini Mitter Co. Inc., Bend, OR) of 54 adults (Reid *et al.*, 2014). Many other studies leveraging wearable light sensor technology have been published (Goulet *et al.*, 2007; Miller *et al.*, 2010; Fields, Linnville and Hoyt, 2016; Dautovich *et al.*, 2019; Ulaganathan *et al.*, 2019).

In many of these studies, participants had to wear the light sensors for an extended period of time. Participants are usually exactly instructed when and how to wear the sensor, and under which circumstances the sensor can be taken off. This currently represents a hurdle to extend data collection efforts to a larger scale. In addition, longer periods of wear time increase the risk of participants’ misuse of the device and/or non-compliance, such as taking off the light sensor and forgetting to put it on again, or accidentally covering it with clothing, resulting in potentially unrepresentative and/or invalid light data. To improve individual exposure estimation, and reduce error, reproducible procedures to exclude invalid data, *i.e.* light data when the participant has not worn the sensor, are necessary. Still, how to best identify invalid light data in an objective and standardized way has, to our knowledge, not been systematically addressed so far.

The determination of invalid data for light is a much harder problem than for example the determination of invalidity of accelerometer-based actigraphy measurements, where a straightforward intuitive approach is to just classify extended periods with no activity counts at all as non-wear time/invalid; the established algorithms for determining non-wear time in actigraphy are indeed based on this idea (Troiano *et al.*, 2008; Hecht *et al.*, 2009; Choi *et al.*, 2011; van Hees *et al.*, 2011, 2013; Syed *et al.*, 2020)). For light, such methods are not useful. At night, during sleep, for example, it is not unusual to have an extended period of no/very low light levels. During computer work, when watching TV or reading in the evening, light intensity levels could be relatively constant, too, and these data are part of an individual’s 24h light exposure profile, and relevant for physiology, as recently reported (Cain *et al.*, 2020). During a typical day, there are usually also periods with spikes and high fluctuation in light intensity, that are due to individuals moving through indoor and outdoor space, and changes in body position. Reliably filtering out short periods of time where the sensor was not worn is therefore hard. However, many physiologically relevant light measures are aggregated over a longer period of time, such as, among many others, average light intensity in lux, variability of the light intensity (quantified for example by standard deviation), the time spent above or below a specific lux threshold, or the timing of light exposure. All these measures are first calculated on a per-day basis, or for periods extending a couple of hours before sleep onset and/or after sleep offset (Reid *et al.*, 2014; Wams *et al.*, 2017), and then averaged over the whole study period. Identifying those days where sensors were not worn throughout the day, and which thus have missing individual-level exposure data despite having recorded light measurements, might therefore be important and might result in reduced measurement error. However, the degree to which invalid light data as typically observed in field studies affects the results has never been systematically investigated.

In this study, we propose a simple algorithm predicting if the sensor-measured light exposure of a given day is potentially biased due to non-compliance (or non-wear time) and should thus be excluded from further analysis. For this purpose, we used data from a study specifically designed to give good training data for our algorithm, where we simulated non-compliance by instructing participants to not wear the light sensor for pre-specified amounts of time, together with data of an ongoing field study where participants’ compliance in wearing the sensor as instructed was high. Light data was captured using the f.luxometer light sensor (f.lux®, Los Angeles), a pendant-worn light wearable recording light in gaze direction. We investigated how much physiologically relevant light metrics, such as average 24h light, average daytime and nighttime light, as well as time spent above specific lux thresholds, differ if calculated based on (i) all available days, (ii) only true valid days (as reported by participants), and (iii) only algorithm-determined valid days. We demonstrate that our algorithm, which is based on an epoch-to-epoch light intensity change metric, performs well in detecting days with extended periods of non-wear time. Furthermore, we demonstrate that light metrics calculated from algorithm selected days approximate calculations based on true valid days well, but also that compared to values of light metrics when calculated based on all light data as it is without exclusions, differences are small.

## MATERIAL AND METHODS

### The f.luxometer light sensor

Throughout our study, f.luxometer (f.lux Software LLC) light sensors were used for continuous measurements of light intensity (**Figure 1**). The f.luxometer is a pendant-worn light sensor that continuously records light using a digital red, green, and blue color light sensor with an infrared-blocking filter (ISL29125, Renesas). The sensor’s sensitivity covers 0.152 to 10,000 lux, with a sampling frequency up to every 10s, with maximum recording times of 2 weeks once fully charged. The f.luxometer has been calibrated to a reference meter (Photoresearch PR-655) and shows high accuracy (error range: 5-25% when tested with 10 widely used commercially available light sources) and precision (5% variability in constant light conditions; between-sensor correlation r=1.0 for 7-day recordings in natural light environments. Sampling frequency was two minutes across all devices.

**Figure 1.**
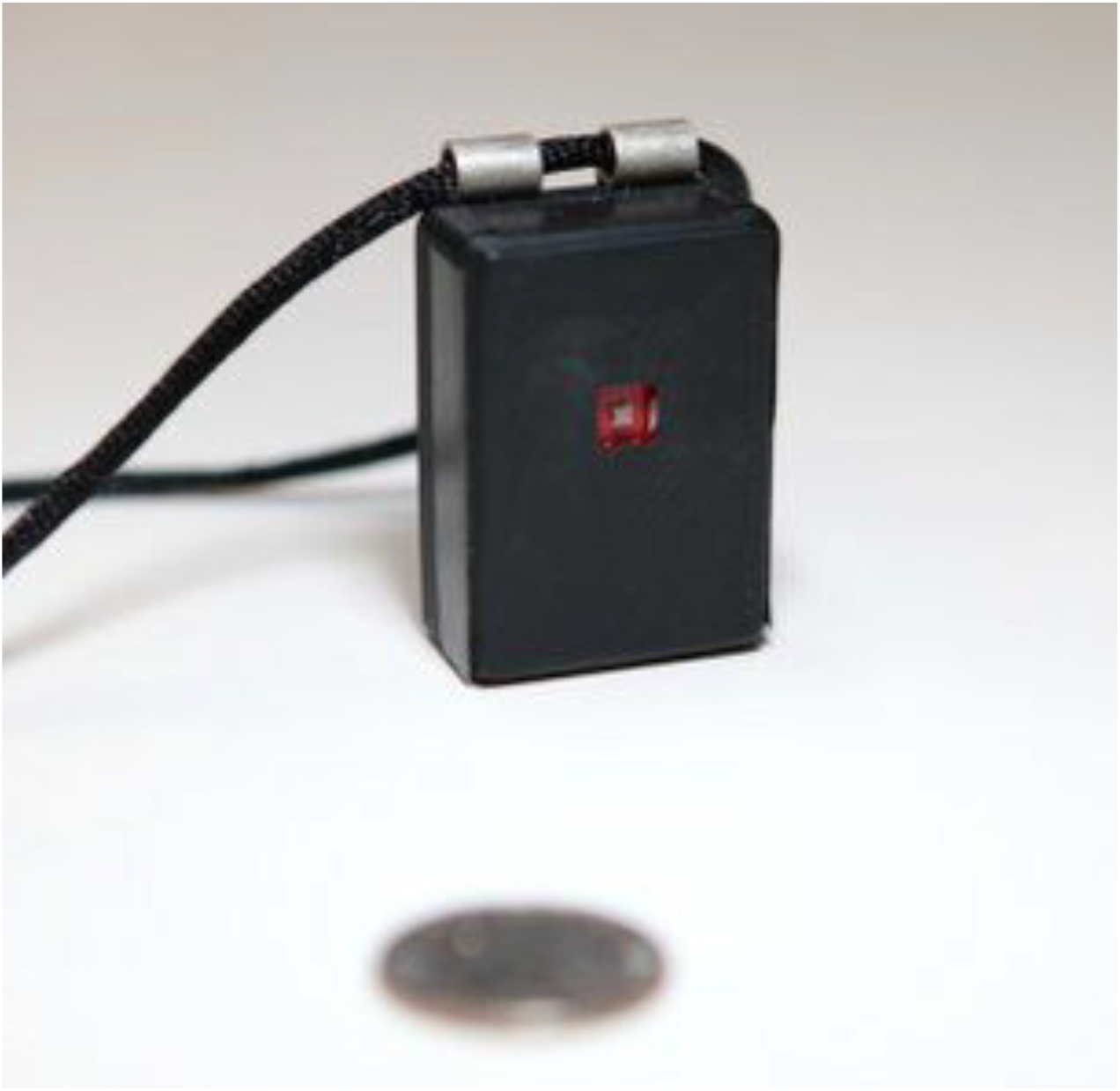
The f.luxometer light sensor.

### Study participants and light data

In our study, we used data specifically designed for the purpose of this study (which was the development of a reliable algorithm for detecting invalid light data days), with an abundance of low-compliance days (“simulated low compliance group”), and data from an ongoing field study where compliance was high (“high compliance group”). This pilot study was approved by the Institutional Review Board (IRB) of the University of Colorado (protocol number 19-0521) and each participant provided informed written consent. The simulation study where low compliance patterns were generated was not considered human subjects research and thus did not require IRB approval. Recruitment of participants and therefore light measurements took place in Boulder, Colorado (latitude 40 1′N, 105 17′W), during the winter time (December 2019 to February 2020) for the high compliance group, and in early summer (June 2020) for the simulation study.

#### High compliance group

This group consisted of four participants of an ongoing pilot study whose main purpose is to assess the impact of an educational light intervention on light profiles, circadian rhythms, sleep, and mood. Participants wore f.luxometer light sensors for 3 weeks (~21 days) each. Participants were instructed to wear the f.luxometer sensors around their necks at all times except for when sleeping, when sensors should be placed near eye level (e.g., on a bedside table). For the duration of the 3-week study period, participants were instructed to keep daily logs to record (i) any f.luxometer sensor removal, and, in case they did remove the sensor, (ii) the time the sensor was removed and the time the sensor was put back on for each day of the study.

#### Simulation study for low compliance

To ensure sufficient data covering the whole spectrum of compliance (from low to high) and, because in the aforementioned field study compliance was in general high, we simulated, in an additional study, low compliance in wearing the f.luxometer sensor in four participants. Participants wore f.luxometer light sensors for 10-14 days, and were instructed to wear sensors around their necks at all times except when sleeping, when sensors were placed near eye level. Participants were also instructed to deliberately remove the sensor for a specified amount of time each day (ranging from 0-24 hours) and maintained logs during the duration of their participation. In these logs, they were asked to record (i) the time the sensor was put on after rising in the morning and the time the sensor was taken off before going to bed in the evening each day, and (ii) the time the sensor was deliberately removed and the time the sensor was put back on during the day each day. Additionally, participants were asked if they followed instructions for wear each day.

### Classification algorithm

For our proposed algorithm, all of the epoch-to-epoch changes in light intensity were transformed on the log-scale (Thapan, Arendt and Skene, 2001; Hut *et al.*, 2008) with a pre-specified offset value (to avoid 0s, which cannot be log-transformed). We then calculated epoch-to-epoch changes *LC_i_(Offs)* per day for the 2-minute epoch length data as:

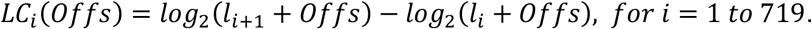

Specifically, *l*_*i*_ are the 720 light intensity measurements per day in temporal order, so that *l*_*i*_ and *l*_*i*+1_ are each two minutes apart, and *Offs* is a real number >0. Our algorithm assigns each day a numerical score, which is defined as the (sample) standard deviation of these 719 change values (in **Figures 2A and 2B** examples for two day with a low and with a high score are presented).

**Figure 2.**
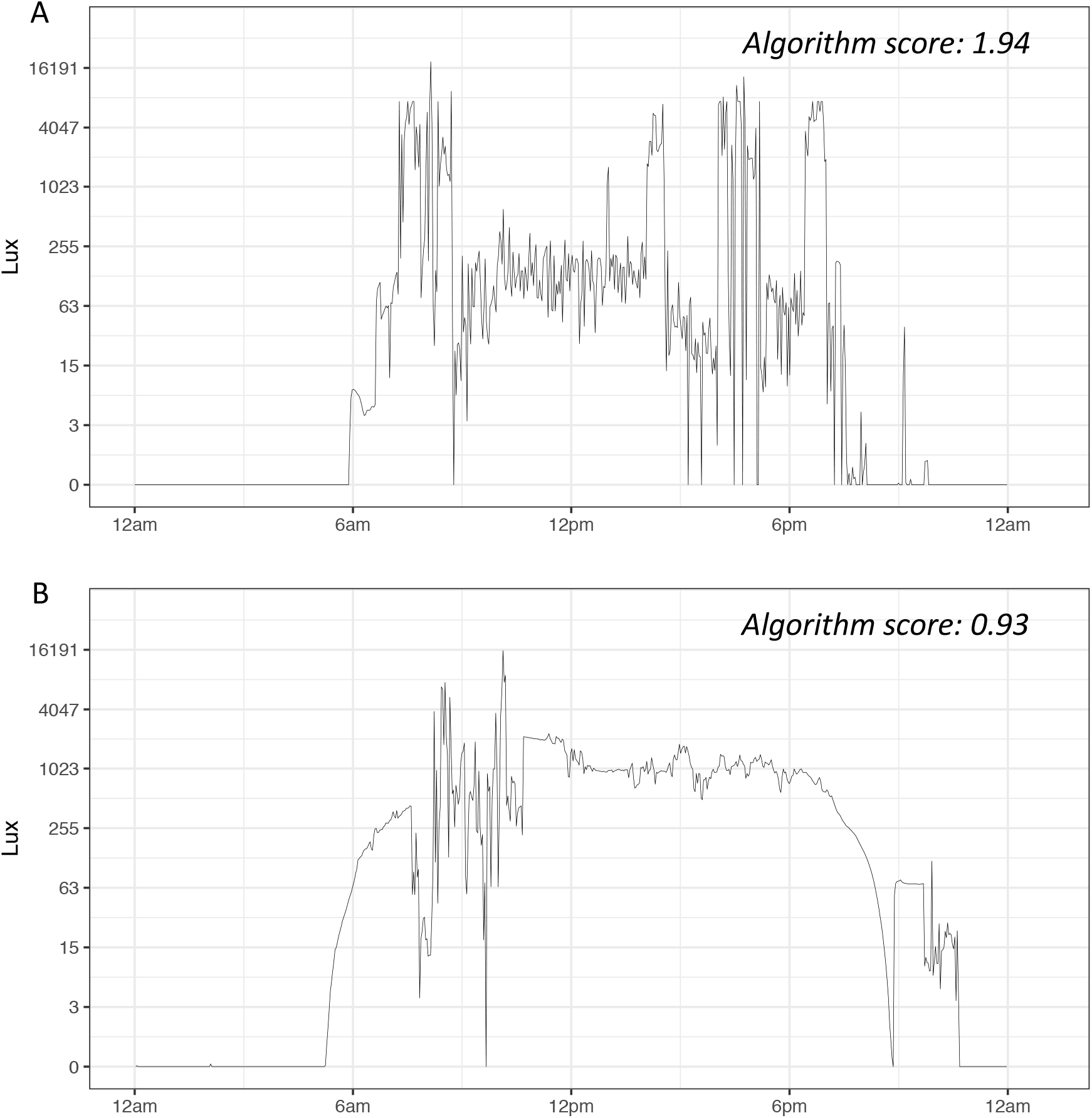
Log-transformed light exposure (lux) across the 24-hour period for (A) a representative valid day of data collection where the sensor was worn with no >4 hour time off and with a calculated algorithm score of 1.94; and (B) a representative invalid day of data collection where the sensore was worn with >4 hour time off and with a calculated algorithm score of 0.93.

Since discriminative performance of our algorithm might also depend on the sampling frequency of the light data, we repeated this approach also by simulating epoch lengths of 4-min, 6-min, 8-min, and 10-min (by taking every 2nd, 3rd, 4th, and 5th epoch). The upper bound for the index *i* changes from 719 to 359, 239, 179, and 143, respectively, otherwise the formulas remain unaltered.

### Statistical analyses

The ability of the score provided by our algorithm to discriminate between valid and invalid days was assessed by receiver operating characteristic (ROC) curves and the corresponding area under the curve (AUC) (also called the c-statistic) (Zou, O’Malley and Mauri, 2007). In addition, we determined the cut-off value maximizing the Youden index (Youden index = sensitivity+(specificity-1)), which is seen as the optimal cut-off value in many applications (Youden, 1950). We also calculated the sensitivity (*i.e.*, the proportion of valid days correctly classified as valid), and the specificity (*i.e.*, the proportion of invalid days correctly classified as invalid) corresponding to this cut-off value. All measures are given with 95% confidence intervals (CIs). Valid days were determined using participant self-reports about light sensor wear times from participants’ logs. Any day in which the participant recorded ≥4 hours of sensor non-wear time during the waking period was marked as invalid. Additionally, the first and last day of recording for each participant were automatically marked as invalid as these days were incomplete and always had ≥4 hours without recording. For our primary analysis, we chose a threshold of 4 hours or more to mark a day as invalid, as this cut-off value has been previously used for determining validity of data from wearables, such as actigraphy (Patel *et al.*, 2015). A missing period of 4h during wake time would also be equivalent to ≥25% missing data, assuming 16h of wake time out of the 24h day. Another prior study has used this criterion for excluding light data as well (Reid *et al.*, 2014). In secondary analyses, we also investigated the performance of the algorithm using alternative definitions of invalidity, specifically ≥2-hour, ≥3-hour, and ≥5-hour non-wear time. Further, to test how much results depended on the specific offset value added to the light data before the log-transformation, we repeated the analyses for various different offset values (specifically 1, 5, 15, 50, and 100). We first performed all analyses with the pooled data across both the high compliance and simulated low compliance groups.

Finally, we report continuous measures of light that are often used in the literature and have been shown to be physiologically important calculated based on all days, self-reported invalid days only, self-reported valid days only, and algorithm based valid days only for all participants pooled (N=8) and stratified by compliance group (N_high compliance_=4, N_low compliance_=4). Specifically, we report (i) average and standard deviation of light exposure across 24 hours (midnight to midnight), (ii) time spent above 50, 100, and 500 lux thresholds, (iii) day-vs. nighttime [7AM-7PM vs. 7PM-7AM] average light intensity (in lux), as well as (iv) time spent above 50 lux thresholds for both day (7AM-7PM) and nighttime (7PM-7AM). For this analysis, we used our main definition of invalidity with a cut-off value of 4h. Specifically, we compared values for each variable based on all days of recording, participant self-reported invalid and valid days. Furthermore, we compared participant self-reported valid days and algorithm based valid days. Algorithm based valid days in this case were determined by our proposed algorithm using the cutoff value in which results of the ROC analyses were optimized. Differences were evaluated using the Mann-Whitney U-test to compare self-reported invalid vs. valid day light metric values, and the partially overlapping samples t-test (Derrick *et al.*, 2017) to compare self-reported valid days vs. algorithm valid days. We also plotted log transformed light exposure (lux) across the 24-hour period including all days vs. participant self-report valid days and participant self-report valid days vs. algorithm determined valid days.

All statistical tests were two-sided at a significance level of 0.05. Analyses were conducted and figures generated in R (R Foundation for Statistical Computing, Vienna, Austria), version 4.0.3.

## RESULTS

### Participant characteristics

We included continuous light recordings from a total of 8 participants (4 in the high compliance group and 4 in the simulated low compliance group) for this analysis. Participants were on average 30.5 years old and 50% were male, with similar distributions among high-compliance (N=4) and low-compliance (N=4) groups. Across all participants, there were 130 days of light recordings, 41 of these days were self-reported by participants as invalid (>4h recorded not worn).

### Algorithm performance

Results from the primary ROC analyses (using a 4-hour threshold for validity and 2-minute epoch length) with offset values of 1, 5, 15, 50, and 100 can be found in **Table 1**. The score given by the algorithm discriminates best between valid and invalid days for an offset of 1 (AUC: 0.77, 95% CI: 0.67-0.87). The optimal cut-off value (*i.e.*, highest Youden index) is 1.36, with sensitivity of 0.87 (95% CI: 0.79-0.93), and specificity of 0.63 (95% CI: 0.49-0.78), respectively. Of the total 130 days of recording included in these analyses, using the cutoff value of 1.36 for the algorithm, 38 days were classified as invalid (26 of these were truly invalid, as reported by participants).

**Table 1.**
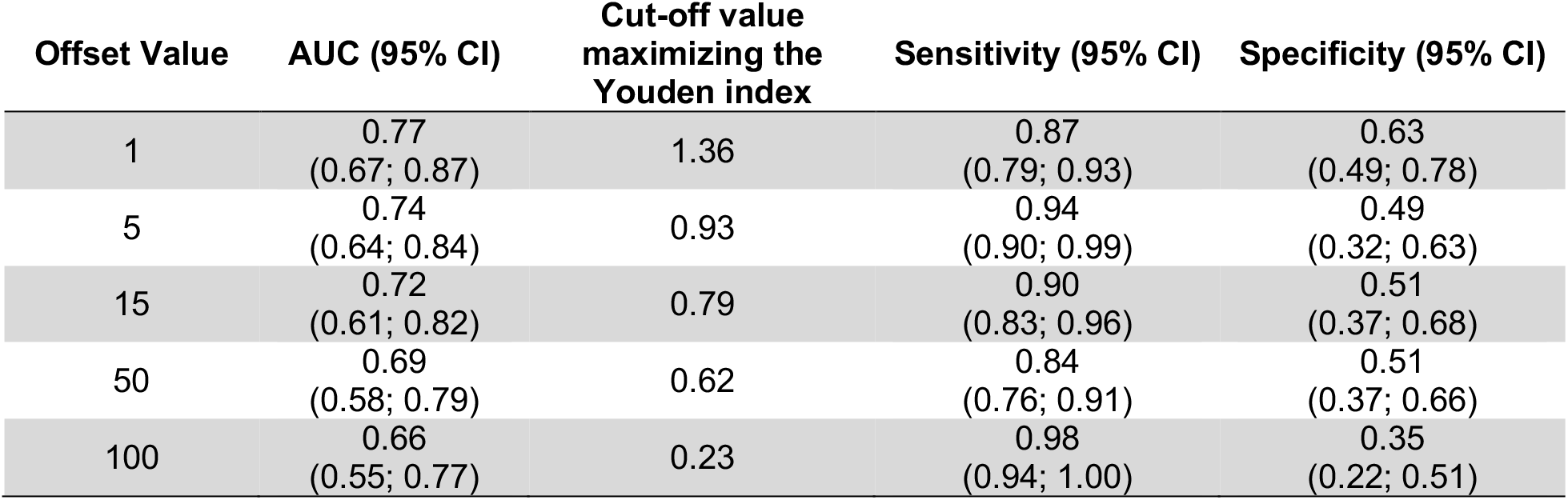
Receiver Operating Characteristic (ROC) analyses results of the Invalid Light Day Classifying Algorithm, with offset values of 1, 5, 15, 50, and 100. Analyses performed in the pooled high and low compliance data with 130 days of light recording. AUC – Area under the receiver operating characteristic curve; CI – confidence interval. Results shown use a 4-hour threshold for validity, and a 2-minute epoch length.

| Offset Value | AUC (95% CI) | Cut-off value<br>maximizing the<br>Youden index | Sensitivity (95% CI) | Specificity (95% CI) |
| --- | --- | --- | --- | --- |
| 1 | 0.77<br>(0.67; 0.87) | 1.36 | 0.87<br>(0.79; 0.93) | 0.63<br>(0.49; 0.78) |
| 5 | 0.74<br>(0.64; 0.84) | 0.93 | 0.94<br>(0.90; 0.99) | 0.49<br>(0.32; 0.63) |
| 15 | 0.72<br>(0.61; 0.82) | 0.79 | 0.90<br>(0.83; 0.96) | 0.51<br>(0.37; 0.68) |
| 50 | 0.69<br>(0.58; 0.79) | 0.62 | 0.84<br>(0.76; 0.91) | 0.51<br>(0.37; 0.66) |
| 100 | 0.66<br>(0.55; 0.77) | 0.23 | 0.98<br>(0.94; 1.00) | 0.35<br>(0.22; 0.51) |

In further analyses, we examined the role of epoch length and of valid day definition on our results, using simulated 4-, 6-, 8-, and 10-minute epoch lengths, and non-wear time of 2-, 3-, and 5-hours for the definition of a valid day. Our results demonstrate that the epoch length does not substantially influence the discriminatory ability of the algorithm. For example, using a 4-hour threshold for validity and 4-minute epoch length, results are as follows: AUC=0.76 (0.66-0.86), sensitivity=0.93 (0.88-0.98), specificity=0.54 (0.39-0.68), and the optimal cut-off value is 1.35 (**Table 2**). These values (except the specific cut-off value) do not vary greatly when compared to results using the same threshold for validity but 10-minute epoch length (AUC: 0.77 (0.68-0.86); sensitivity: 0.92 (0.87-0.97); specificity: 0.56 (0.41-0.71); optimal cut-off value: 1.60; Table 2). On the other hand, the criterion for validity does influence the discriminatory ability of the algorithm. Using a 2-minute epoch length, the algorithm performs better with a 5-hour threshold for validity (AUC: 0.85 (0.76-0.93); sensitivity: 0.94 (0.89-0.98); specificity: 0.65 (0.47-0.79); optimal cut-off value: 1.22) in comparison to a 2-hour threshold for validity (AUC: 0.68 (0.58-0.78); sensitivity: 0.94 (0.87-0.99); specificity: 0.45 (0.32-0.58); optimal cut-off value: 1.24; **Table 2**).

**Table 2.**
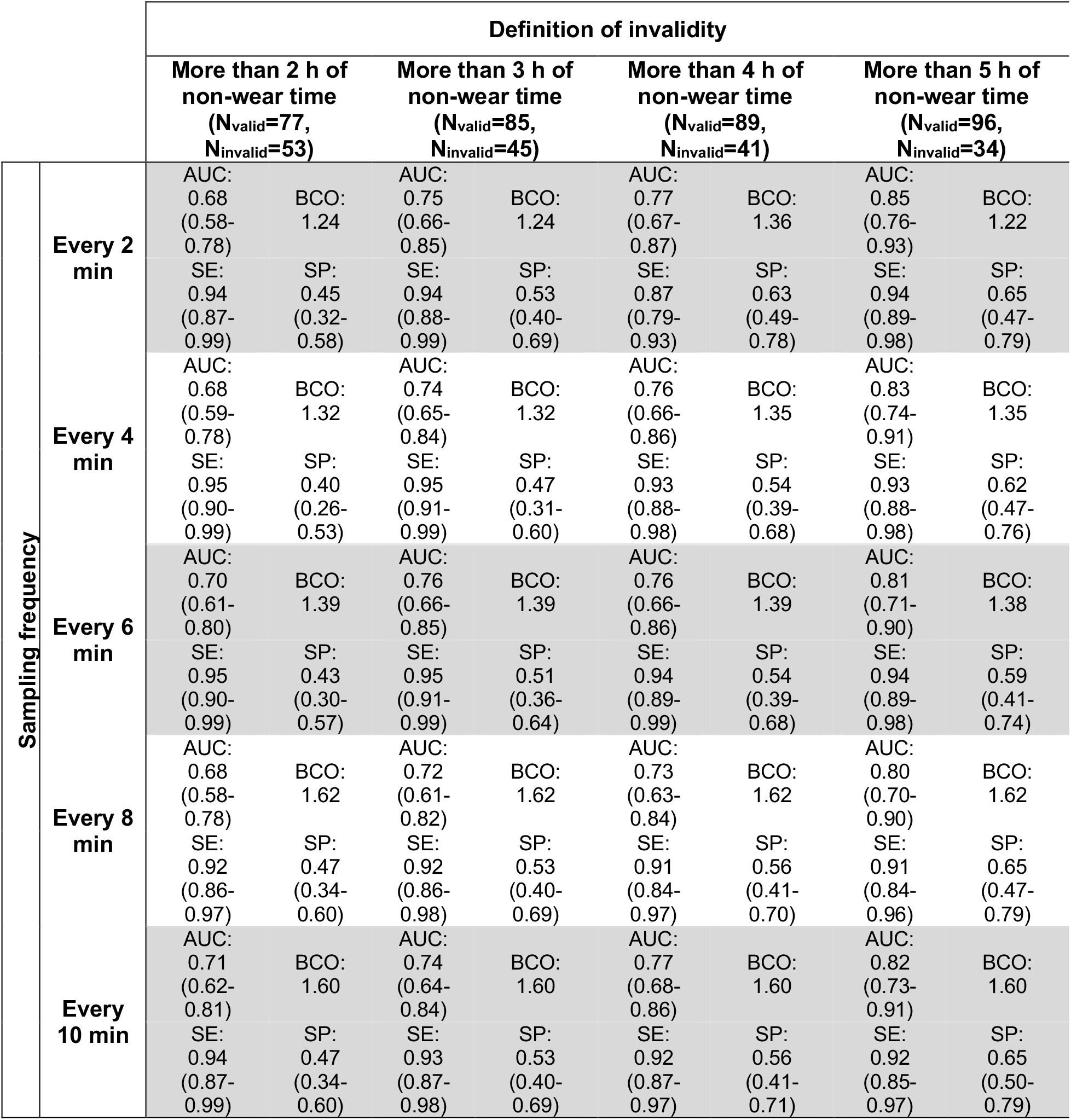
Performance of the Invalid Light Day Classifying Algorithm in detecting invalid days by sampling frequency and by the chosen definition of invalidity (i.e. up to which amount of non-wear time a day is still considered valid) for the pooled high and low compliance data with 130 days of light recording. AUC – Area under the receiver operating characteristic curve; BCO – best cut-off value (i.e., maximal Youden index); SE – Sensitivity; SP – Specificity.

### Light exposure profiles

We compared computed light variables when calculated based on all 130 days included in the study, based on the 89 valid and 41 invalid days according to participant self-report (using a >4-hour threshold), and based on the 92 days which were rated as valid according to the algorithm (*i.e.*, days that had an algorithm score of ≥1.36) (**Table 3**). Results in most aggregated light metrics were similar when calculated for algorithm-determined valid days vs. self-reported valid days. An exception was the metric “variability (standard deviation) in 24h lux” which was significantly overestimated in the algorithm based valid days compared to the self-reported valid days (median_algorithm valid days_=974.9 vs. median_self-reported valid days_=760.6). Time spent above a 500-lux threshold in minutes (p=0.05), and average daytime (7AM-7PM) light exposure in lux (p=0.02) were also slightly increased when calculated based on the algorithm valid days (**Table 3**). Mean light exposure across the 24-hour period including algorithm determined valid days vs. participant self-report valid days only did not differ overall (median_algorithm valid days_=264.3 vs. median_self-reported valid days_=200.5, p=0.14; **Table 3**) nor at any time point (**Figure 3A**).

**Table 3.** Descriptive statistics of basic light exposure metrics calculated based on all days of recording, participant self-reported invalid and valid days of recording only, and algorithm valid days only (days that had a score of ≥1.36 according to the algorithm). Values are presented as median (interquartile range). Significant p-values are marked in bold (p<0.05)

| Light Metric | All Days<br>(N=130) | Self-Report<br>Invalid Days<br>(N=41) | Self-Report<br>Valid Days<br>(N=89) | Algorithm<br>Valid Days<br>(N=93) | P-value<br>(Self-report<br>invalid vs.<br>valid)<sup>1</sup> | P-value<br>(Self-report<br>valid vs.<br>algorithm<br>valid)<sup>2</sup> |
| --- | --- | --- | --- | --- | --- | --- |
| Algorithm Score | 1.56<br>(1.28; 1.74) | 1.16<br>(0.24; 1.57) | 1.64<br>(1.42; 1.84) | 1.68<br>(1.53; 1.88) |  |  |
| Average 24h light<br>(lux) | 212.5<br>(122.9; 404.4) | 300.4<br>(113.8; 449.0) | 200.5<br>(131.2; 372.7) | 264.3<br>(153.6; 420.0) | 0.38 | 0.14 |
| Variability (standard<br>deviation) of 24h lux | 801.2<br>(421.0;<br>1347.6) | 855.2<br>(267.3; 1356.7) | 760.6<br>(451.9; 1287.4) | 974.9<br>(616.3; 1438.7) | 0.96 | **<0.001** |
| Time spent above 50-lux threshold<br>(minutes) | 489.0<br>(347.5; 593.0) | 340.0<br>(234.0; 662.0) | 510.0<br>(426.0; 584.0) | 496.0<br>(404.0; 582.0) | 0.09 | 0.43 |
| Time spent above<br>100-lux threshold<br>(minutes) | 353.0<br>(200.5; 476.5) | 232.0<br>(96.0; 522.0) | 388.0<br>(286.0; 470.0) | 362.0<br>(266.0; 462.0) | **0.04** | 0.67 |
| Time spent above<br>500-lux threshold<br>(minutes) | 104.0<br>(440.0; 160.5) | 104.0<br>(34.0; 198.0) | 100.0<br>(50.0; 154.0) | 110.0<br>(66.0; 164.0) | 0.90 | **0.05** |
| Average daytime<br>(7AM-7PM) light (lux) | 411.8<br>(237.9; 788.4) | 598.2<br>(237.2; 933.8) | 380.7<br>(246.5; 730.4) | 503.5<br>(292.1; 805.7) | 0.26 | **0.02** |
| Daytime (7AM-7PM)<br>time spent above 50-lux threshold<br>(minutes) | 454.0<br>(332.0; 552.0) | 328.0<br>(194.0; 608.0) | 478.0<br>(382.0; 544.0) | 460.0<br>(362.0; 544.0) | 0.14 | 0.28 |
| Average nighttime<br>(7PM-7AM) light (lux) | 11.1<br>(4.5; 19.6) | 11.2<br>(4.6; 33.2) | 11.0<br>(4.5; 16.5) | 11.2<br>(6.0; 19.6) | 0.29 | 0.06 |
| Nighttime (7PM-7AM)<br>time spent above 50-lux threshold<br>(minutes) | 26.0<br>(10.0; 49.5) | 24.0<br>(8.0; 50.0) | 26.0<br>(10.0; 48.0) | 28.0<br>(12.0; 50.0) | 0.66 | 0.64 |

<sup>1</sup>p-values from Mann-Whitney U-tests.
<sup>2</sup>p-values from partially overlapping samples t-tests (Derrick *et al.*, 2017).

**Figure 3.**
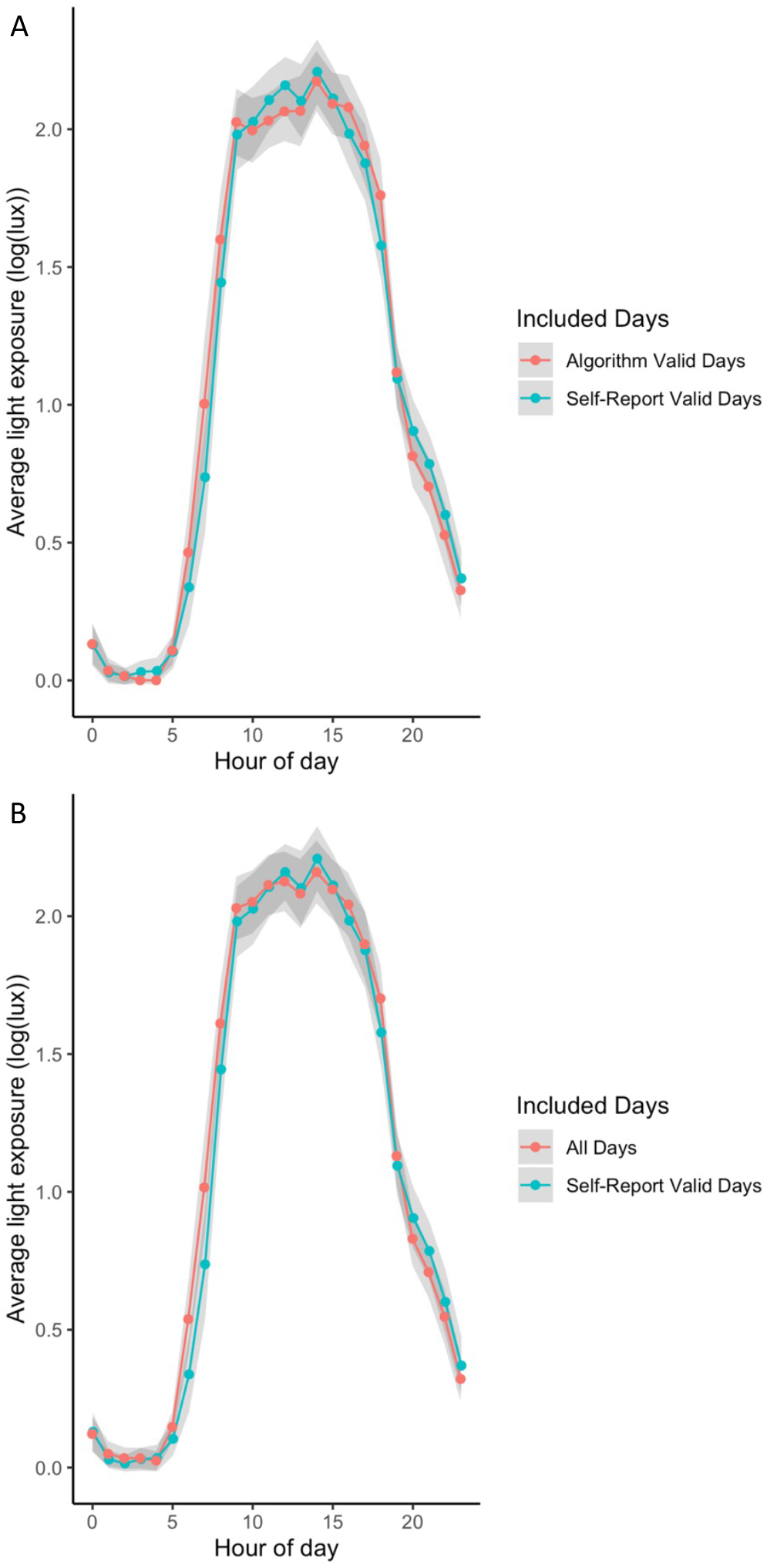
Mean values of log-transformed light exposure (lux) across the 24-hour period including (A) algorithm determined valid days (N=93, days that had a score of ≥1.36 according to the algorithm) (red) compared to participant self-reported valid days (N=89, using a >4-hour threshold) (blue); and (B) all days of data collection (N=130) (red) compared to participant self-reported valid days (N=89, using a >4-hour threshold) (blue). Grey shading indicates 95% confidence intervals of calculated values.

Additionally, the values of the light measurements did not differ when including all days (N=130) as compared to only participant self-reported valid days (N=89), except for the amount of time spent above 100-lux (min) (median_all days_=353 minutes vs. median_self-reported valid days_=388 minutes, p=0.04, **Table 3**). Light exposure across the 24-hour period including all days vs. participant self-report valid days only did not differ overall (p=0.38; **Table 3**) nor at any time point (**Figure 3B**). Among the four participants of our high compliance group, values of the light metrics did not vary to a large extent when calculated based on all days, self-reported valid days only, and algorithm valid days only (**Table 4**). However, among the four participants of our simulated low compliance study, using algorithm valid days only for the calculation of some light metrics better reflected values obtained using participant self-report valid days only compared to using all days of data collection (**Table 4**). Despite marked differences in some of the light metrics amongst the high compliance group vs. the low compliance group (e.g. median_average 24h light (lux), high compliance_ = 163.3 (Q1: 114.5, Q3: 276.7), median_average 24h light (lux), low compliance_ = 403.8 (Q1: 235.2, Q3: 526.5)), algorithm calculated scores were in comparable ranges for both of these groups (High compliance = 1.54 (Q1: 1.34, Q3: 1.73), Low compliance = 1.45 (Q1: 1.16, Q3: 1.89)).

**Table 4.**
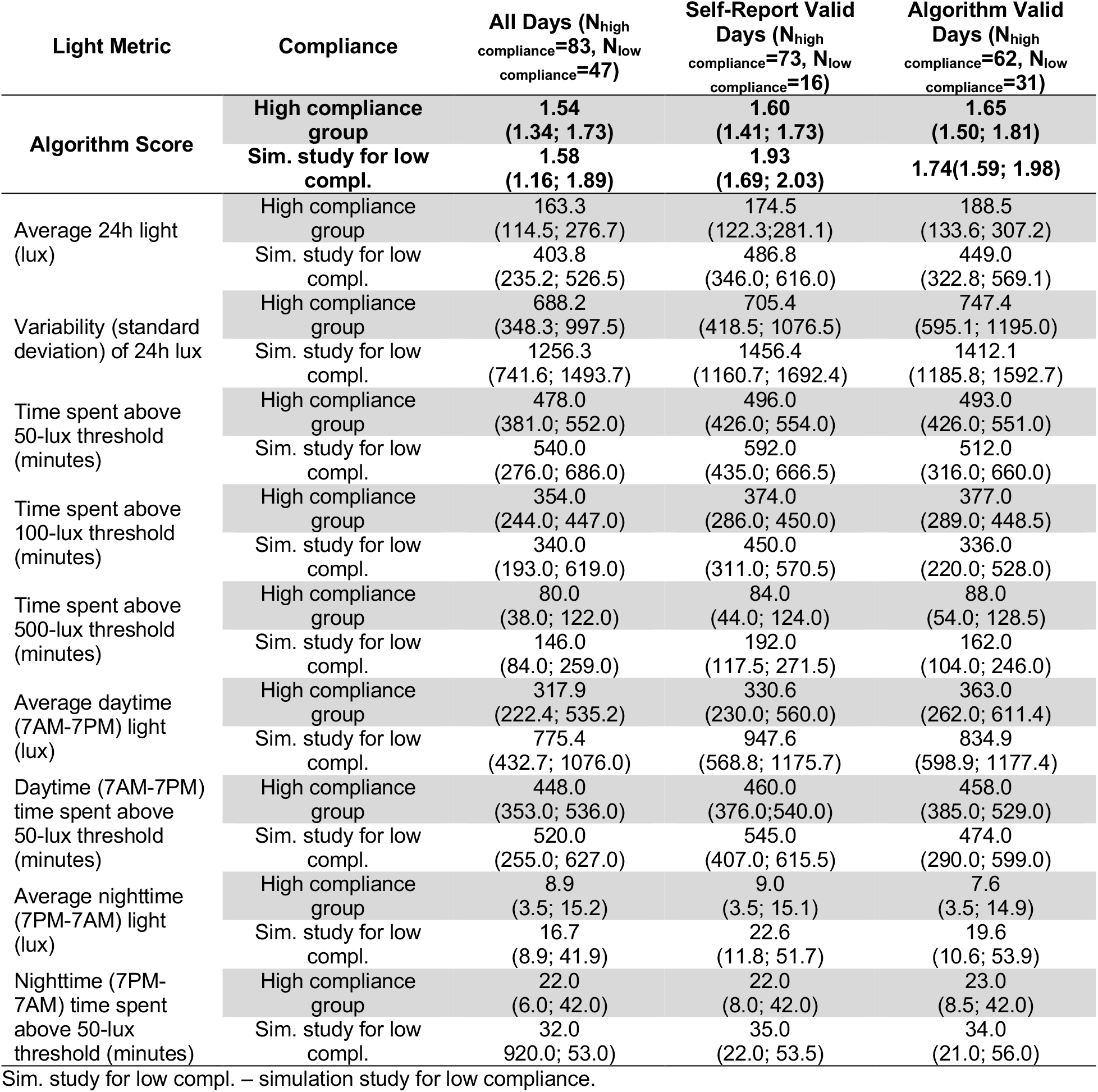
Descriptive statistics of basic light exposure metrics calculated based on all days of recording, participant self-reported invalid and valid days of recording only, and algorithm valid days only (days that had a score of ≥1.36 according to the algorithm) *stratified by compliance*. Values are presented as median (interquartile range).

| Light Metric | Compliance | All Days ($N_{\text{high compliance}}=83$, $N_{\text{low compliance}}=47$) | Self-Report Valid Days ($N_{\text{high compliance}}=73$, $N_{\text{low compliance}}=16$) | Algorithm Valid Days ($N_{\text{high compliance}}=62$, $N_{\text{low compliance}}=31$) |
| --- | --- | --- | --- | --- |
| Algorithm Score | High compliance group | 1.54<br>(1.34; 1.73) | 1.60<br>(1.41; 1.73) | 1.65<br>(1.50; 1.81) |
|  | Sim. study for low compl. | 1.58<br>(1.16; 1.89) | 1.93<br>(1.69; 2.03) | 1.74(1.59; 1.98) |
| Average 24h light (lux) | High compliance group | 163.3<br>(114.5; 276.7) | 174.5<br>(122.3; 281.1) | 188.5<br>(133.6; 307.2) |
|  | Sim. study for low compl. | 403.8<br>(235.2; 526.5) | 486.8<br>(346.0; 616.0) | 449.0<br>(322.8; 569.1) |
| Variability (standard deviation) of 24h lux | High compliance group | 688.2<br>(348.3; 997.5) | 705.4<br>(418.5; 1076.5) | 747.4<br>(595.1; 1195.0) |
|  | Sim. study for low compl. | 1256.3<br>(741.6; 1493.7) | 1456.4<br>(1160.7; 1692.4) | 1412.1<br>(1185.8; 1592.7) |
| Time spent above 50-lux threshold (minutes) | High compliance group | 478.0<br>(381.0; 552.0) | 496.0<br>(426.0; 554.0) | 493.0<br>(426.0; 551.0) |
|  | Sim. study for low compl. | 540.0<br>(276.0; 686.0) | 592.0<br>(435.0; 666.5) | 512.0<br>(316.0; 660.0) |
| Time spent above 100-lux threshold (minutes) | High compliance group | 354.0<br>(244.0; 447.0) | 374.0<br>(286.0; 450.0) | 377.0<br>(289.0; 448.5) |
|  | Sim. study for low compl. | 340.0<br>(193.0; 619.0) | 450.0<br>(311.0; 570.5) | 336.0<br>(220.0; 528.0) |
| Time spent above 500-lux threshold (minutes) | High compliance group | 80.0<br>(38.0; 122.0) | 84.0<br>(44.0; 124.0) | 88.0<br>(54.0; 128.5) |
|  | Sim. study for low compl. | 146.0<br>(84.0; 259.0) | 192.0<br>(117.5; 271.5) | 162.0<br>(104.0; 246.0) |
| Average daytime (7AM-7PM) light (lux) | High compliance group | 317.9<br>(222.4; 535.2) | 330.6<br>(230.0; 560.0) | 363.0<br>(262.0; 611.4) |
|  | Sim. study for low compl. | 775.4<br>(432.7; 1076.0) | 947.6<br>(568.8; 1175.7) | 834.9<br>(598.9; 1177.4) |
| Daytime (7AM-7PM) time spent above 50-lux threshold (minutes) | High compliance group | 448.0<br>(353.0; 536.0) | 460.0<br>(376.0; 540.0) | 458.0<br>(385.0; 529.0) |
|  | Sim. study for low compl. | 520.0<br>(255.0; 627.0) | 545.0<br>(407.0; 615.5) | 474.0<br>(290.0; 599.0) |
| Average nighttime (7PM-7AM) light (lux) | High compliance group | 8.9<br>(3.5; 15.2) | 9.0<br>(3.5; 15.1) | 7.6<br>(3.5; 14.9) |
|  | Sim. study for low compl. | 16.7<br>(8.9; 41.9) | 22.6<br>(11.8; 51.7) | 19.6<br>(10.6; 53.9) |
| Nighttime (7PM-7AM) time spent above 50-lux threshold (minutes) | High compliance group | 22.0<br>(6.0; 42.0) | 22.0<br>(8.0; 42.0) | 23.0<br>(8.5; 42.0) |
|  | Sim. study for low compl. | 32.0<br>(920.0; 53.0) | 35.0<br>(22.0; 53.5) | 34.0<br>(21.0; 56.0) |

Sim. study for low compl. – simulation study for low compliance.

## DISCUSSION

The goal of this study was to develop an easy-to-implement algorithm for automated detection of days on which non-wear time exceeds pre-specified thresholds. Systematically evaluating our proposed algorithm’s performance, we showed that 79% of all days in our data were correctly classified as valid/invalid (as assessed via self-reports; AUC=0.77, 95% CI: 0.67; 0.87); and that the discriminatory performance of our automated algorithm, which does not require any participant input, is good enough to give similar values in the derived light variables when excluding days with prolonged periods of non-wear time based on the algorithm’s decision as compared to participant self-reports (with the exception of variability of 24h lux). Therefore, automated detection of valid light data days appears feasible.

However, we also demonstrated that the impact of non-wear time (as determined from diary self-reports) of a pendant-worn light sensor on derived physiologically relevant light variables may not be substantial, especially in high-compliance cohorts. Specifically, when using measures of light exposure which are averaged across multiple days of data collection, and not calculated on a per-day basis, differences between values calculated based on all available data vs. based on only self-reported valid days were small. However, these differences between values grew when data were mainly from the simulated low compliance group where we intentionally generated a larger variety of non-wear time patterns in the data by having participants remove their sensors for specific amounts of time each day. The data from participants in the ongoing field study with a high compliance appeared to have lower levels of differences between light across the board (**Table 4**). Of note, values of some of the calculated light metrics (e.g. average 24h light, variability of 24h lux) markedly differed between the four participants in the high compliance group vs. the four participants in the low compliance group. These differences might incidate high inter-individual variability, probably due to different lifestyle, working, and sleeping habits. Still, our algorithm calculated scores were in comparable ranges for both groups, indicating that the scores are quite insensitive to inter-individual variability, a desireable property for an algorithm to be universially applied. Taken together, this suggests that in studies where compliance is known to be high and light is averaged across multiple days, filtering out invalid days may not be entirely necessary. On the other hand, our algorithm is simple to implement, performs well and provides similar and valid results regarding physiologically important aggregated light metrics (except variability of 24h lux). When calculating light metrics on a per-day basis, careful consideration of invalid days is probably more important, and our algorithmic approach might be superior to an analysis of all data as it is and also preferable to the burden of tracking compliance (with the associated risk of false reports) from the participants via questionnaires.

For our algorithm, we calculated differences in log-transformed light intensity values for all consecutive time-points and then took the per-day SD. The idea was that when the light sensor is stationary located during non-wear time, changes in light intensity should not be abrupt, but rather gradual, if at all, and consequently the calculated SD should get smaller the longer the non-wear time is. By taking the logarithm of the light intensity values we implicitly encode that circadian responses to light follow a log-linear relationship (Thapan, Arendt and Skene, 2001; Hut *et al.*, 2008), and, importantly, ensure that calculated differences do not get too large and hence do not have a disproportionately high impact on the SD. Finally, by taking differences from time-point to time-point, the average light intensity level should be roughly cancelled out, so that our algorithm should be rather unaffected by seasonal or locational differences, although this cannot be directly assessed in our data (only 8 participants, all from the Boulder area) and still has to be validated in future studies. Because the score of our algorithm which determines a day’s validity of light data is based on the SD of modified light lux values, there is a selection towards higher light variability in the algorithm selected valid day. This is the reason why the variability of 24h lux metric is off when using only algorithm selected valid days.

To date, to the best of our knowledge, systematic investigations about invalid data from wearable light sensors and examinations how to best deal with the problem of invalid light data are lacking. Reviewing approaches for data preparation and/or exclusions in previous studies leveraging light sensor data, we found approaches to be very diverse, making comparisons of results difficult. Some studies did not report on exclusions of light data at all and appear to have used the light data for analysis as it was (Papantoniou *et al.*, 2014; Reid *et al.*, 2014; Wams *et al.*, 2017). In two studies, all light measurements of <1 lux during the (self-reported) out-of-bed period were excluded from further analysis (Scheuermaier, Laffan and Duffy, 2010; Obayashi *et al.*, 2014). The authors argue that this way periods where the (wrist-worn) sensor was accidentally covered with a sleeve were excluded. This is a valid argument, however, invalidity includes more scenarios than a sensor covered by clothes, and this approach does not address the issue of non-wear time. In another study, validity of the light data (measured via the AW-L Actiwatch device) was linked to non-zero activity counts, complementing diary information about wear and non-wear times (Goulet *et al.*, 2007), but this approach requires the light sensor to be linked to an activity-count device. Miller et al. leveraged activity and temperature data, which is captured together with light data by the Daysimeter, for identifying noncompliance based on a visual inspection. Partly visual inspections of data have been used to in- or exclude light data also elsewhere (Daugaard *et al.*, 2017; Ulaganathan *et al.*, 2019), but the obvious lack of objective criteria and thus transparency is, beside staff time and cost, a severe drawback of such an approach. Related to this, in a side project to this study, we asked two experts in the field of light and chronobiology to blindly assess a subset of our light data for validity and gave them detailed instruction on which criteria to look at. Still, inter-rater agreement between the two experts was surprisingly low (Cohen’s kappa=0.74), underlining that it is hard to find consensus on valid/invalid days even for experts. **Figure 2** should also be interpreted in this light – although one valid and one invalid day of light data collection is presented, and the selected patterns are somewhat typical, in practice there are various shades and also distinctly different patterns may arise. Finally, information on participants’ light sensor wear and non-wear time was also assessed via self-reports (Goulet *et al.*, 2007), and we used this information (even as the “gold standard”) in our study, too. Albeit a valid approach, self-reports rely on participant compliance, which may be variable depending of the study population, and may be incomplete or erroneous. Especially in multi-outcome field studies with many different wearables, study visits, and questionnaires, reducing participant burden by eliminating additional reports without compromising data quality is asked for, as this in turn likely increases participant motivation and compliance. Our proposal of algorithmically processing light data validity is a promising attempt to circumvent or at least mitigate all of these problems.

We recognize that our methodology has some limitations. First, in our primary analysis, we declared a day to be invalid if the self-reported non-wear time of the sensor exceeds 4h. A missing period of 4h during wake time would translate to 25% missing data, assuming 16h of wake time out of the 24h day. This definition of invalid days has been used previously (Reid *et al.*, 2014), but further exploration is required to see if, depending on the research question, this definition is justified or has to be optimized. However, given the complete absence of guidance at this point, we believe this is a sensible approach. In secondary analyses, we varied the threshold of non-wear time above which days were classified as invalid. As expected, the algorithm identified invalid days the more accurately the higher this threshold was. Second, we based the decision on validity of days on self-reported non-wear time. However, self-reports are error-prone in itself and are therefore not on optimal benchmark to be compared against (*i.e.*, they do not constitute a perfect “gold-standard”). Another drawback of self-reports is that periods of time where the sensor was inadvertently covered by clothes are not captured. This might be less of an issue in the simulation study of low compliance because participants there were well aware that the purpose of them wearing the light sensor was to simulate low compliance, and that for this it was essential to accurately following the instructions about wearing/not wearing and not covering the sensor. The comparison of light variables derived based on self-reported valid days and algorithm determined valid days, as displayed in **Table 3**, should also be viewed in this light, namely that using self-reported valid days for calculation does not necessarily give the “correct” values and deviations between the two variants do not necessarily mean that the variables derived from algorithm-determined valid days are “wrong”. Third, our light data was recorded every 2 minutes, only a moderate resolution compared to other studies were light was recorded in intervals as low as 15 seconds (Papantoniou *et al.*, 2014). However, in a study investigating how measurement duration and frequency of light data affected the obtained data, it was shown that a measurement duration of at least one week and a measurement frequency of two minutes or finer provides the most reliable estimates of personal light exposure measures (Ulaganathan *et al.*, 2017). So, the 2-min gaps between light recordings in our study should be sufficient, and the long follow-up of our participants of about 2 weeks is a strength. On a similar note, the best cut-off value discriminating between valid and invalid days strongly depends on the sampling frequency, as demonstrated in **Table 2**, where cut-off values for 2-min to 10-min gaps are given. Cut-off values for less than 2-min remain to be determined. Fourth, our light data of 130 days comes from 8 young, healthy participants from the Boulder area, known for its many sunny days and inhabitants’ active lifestyle. More work is needed to investigate the performance of our algorithm in other or more diverse populations (e.g. regarding age, health status, pre-existing medical conditions, latitude at which individuals are living), and our approach should be validated in other studies with larger number of participants in various free-living settings. Finally, in our study, we used the wrist-worn f.luxometer as the wearable sensor assessing light, and light was recorded in gaze direction, which is most relevant to humans when studying the physiological and behavioral effects of light (Skene and Arendt, 2006). Investigating how the performance of our approach depends on the specific light sensor used and the location of the light measurement (e.g. wrist, chest, eye-level) would be important as well. Because our algorithm uses modifiable arguments (offset value [set to 1 for our population], and cut-off value [set to 1.36 for our population and 2 minute recording gaps], it could be easily modified and optimized for each study’s needs and adopted to other accelerometers.

In conclusion, our results suggest that our proposed algorithm discriminates well enough between valid and invalid days to minimize distortions in individual-level environmental light recordings and to provide similar light exposure patterns as those that would be generated if using participant self-reports of compliance. We did not see the necessity of filtering out invalid days of collection in the first place, as light levels do not differ greatly between all days and self-report valid days only and compliance in participants was quite high. Still, our algorithm is simple to implement and performs well regardless of the compliance level of the data without requiring participants and researchers to monitor during the study duration. Hence, self-reports on non-wear time, which in themselves are also prone to faulty entries, are dispensable, and can be substituted by application of our algorithm, without compromising the validity of derived light variables. Large-scale field studies and/or clinical trials could likely benefit from implementing our, or a similar, algorithmic approach for automatic detection of days with invalid light data.

## FUNDING

J.F. was funded by a postdoctoral research exchange grant from the Max Kade Foundation, Inc. Otherwise, this research received no specific grant from any funding agency in the public, commercial, or not-for-profit sectors.

## CONFLICT OF INTEREST STATEMENT

Michael Herf is a co-Founder of f.lux Software LLC, which made the prototype sensors used in this study.

